# Effects of Reproductive Experience on Cost-Benefit Decision Making in Females and Males

**DOI:** 10.1101/2023.11.03.565418

**Authors:** Mojdeh Faraji, Omar A. Viera-Resto, Barry Setlow, Jennifer L. Bizon

## Abstract

Almost all individuals undergo reproductive and/or parenting experience at some point in their lives, and pregnancy and childbirth in particular are associated with alterations in the prevalence of several psychiatric disorders. Research in rodents shows that maternal experience affects spatial learning and other aspects of hippocampal function. In contrast, there has been little work in animal models concerning how reproductive experience affects cost-benefit decision making, despite the relevance of this aspect of cognition for psychiatric disorders. To begin to address this issue, reproductively experienced (RE) and reproductively naïve (RN) female and male Long-Evans rats were tested across multiple tasks that assess different forms of cost-benefit decision making. In a risky decision-making task, in which rats chose between a small, safe food reward and a large food reward accompanied by variable probabilities of punishment, RE and RN males did not differ, whereas RE females chose the large risky reward significantly more frequently than RN females (greater risk taking). In an intertemporal choice task, in which rats chose between a small, immediate food reward and a large food reward delivered after a variable delay period, RE males chose the large reward more frequently than RN males, whereas RE females chose the large reward less frequently than RN females. Together, these results show distinct effects of reproductive experience on different forms of cost-benefit decision making in rats of both sexes, and highlight reproductive status as a variable that could influence aspects of cognition relevant for psychiatric disorders.

## 1. Introduction

Reproductive experience (i.e., mating, pregnancy, childbirth, and parenting) is associated with a range of somatic, hormonal, and neural adaptations that prepare organisms to create and raise their offspring (Bhatia & Chhabra, 2018; Hoyt & Falconi, 2015; Johnson & Cipolla, 2015; Mastorakos & Ilias, 2000; Mizuno et al, 2017; Reyes et al, 2018; Tal & Taylor, 2000; Tan & Tan, 2013; Wen et al, 2019). The female mammalian brain in particular undergoes remarkable alterations during the peripartum period. Studies in rodents show shifts in corticotropin-releasing hormone (CRH), opioid, oxytocin, and dopamine signaling, as well as changes in gene expression, along with altered progesterone and estrogen profiles during the peripartum period (Alcántara-Alonso et al, 2017; Brunton, 2019; Brunton & Russell, 2008; Cárdenas et al, 2020; Kinsley & Amory-Meyer, 2011; Robinson et al, 2011; Shnitko et al, 2017). In humans, there are changes in neural organization and resting state brain activity (Hoekzema et al, 2022), as well as alterations in grey matter volume and cortical thickness during pregnancy (Rehbein et al, 2022) and early postpartum (Chechko et al, 2022; Servin-Barthet et al, 2023). Many of these brain changes (particularly those observed in rodents) have been linked to offspring-relevant behaviors, including gestation, birth, lactation, pair bonding/affiliation, and maternal aggression (Arévalo & Campbell, 2020; Hahn-Holbrook et al, 2011; Leuner et al, 2010; Walter et al, 2021). The peripartum period is also associated with changes in vulnerability to psychiatric disorders, however, including depression (Barba-Müller et al, 2019; Morgan et al, 2021; Osborne et al, 2022; Pawluski et al, 2017; Skalkidou et al, 2012; Worthen & Beurel, 2022), anxiety, puerperal psychosis, obsessive-compulsive disorder, and post-traumatic stress disorder (Barba-Müller et al, 2019; Meltzer-Brody et al, 2018; Miller et al, 2015a; b). The causes of such vulnerability changes are unclear and likely multifaceted (e.g., genetic or environmental predispositions, hormonal fluctuations, peripartum shifts in socioeconomic conditions). As such, the use of animal models can provide experimental control that can help to identify biological and/or environmental contributions to the effects of reproductive experience on behavior.

Despite numerous studies documenting links between reproductive experience and changes in risks of psychiatric disorders, most preclinical research on the consequences of reproductive experience (outside that tied explicitly to offspring-directed behaviors) has focused on the hippocampus and hippocampal functions such as learning and memory (Eid et al, 2019; Maeng & Shors, 2012; Moses-Kolko et al, 2021; Pawluski et al, 2006; Roes & Galea, 2016; Shors, 2016). In contrast, other brain systems implicated in psychiatric disorders (e.g., prefrontal cortex and amygdala) and associated aspects of cognition (e.g., executive functions, including decision making) have received less attention. In humans, both increased (Kim et al, 2018) and decreased (Hoekzema et al, 2017) grey matter volume and thickness of prefrontal cortex have been reported following pregnancy and childbirth. In addition, the default mode network (DMN), a closely interconnected system of brain regions (including prefrontal cortex) representing the brain’s baseline activity that is involved in self-perception and social cognition, is functionally altered during pregnancy and after childbirth. Hypoactivity in the DMN during working memory (Bak et al, 2020) and altered coherence of DMN across pregnancy in response to infant cues are examples of such changes (Hoekzema et al, 2022). Alterations in decision making are prevalent in a range of psychiatric disorders, including those linked to reproductive experience (Cáceda et al, 2014; Morris, 2018; Mukherjee et al, 2020; Purcell et al, 2022; Teng et al, 2016; Valyan et al, 2020). As such, a better understanding of the effects of reproductive experience on executive functions/decision making could lead to new insights into the etiology of peripartum changes in risks of psychiatric disorders.

The majority of research on the effects of reproductive experience on brain and behavior has focused on the peripartum period (i.e., during pregnancy and/or early child rearing). A growing body of evidence, however, suggests that reproductive experience can have effects that far outlast this period (Duarte-Guterman et al, 2019; Hoekzema et al, 2017). For example, a neuroimaging study in humans revealed reductions in grey matter in a cohort of primiparous mothers that persisted for six years postpartum (Martínez-García et al, 2021), and in another study, grey matter volume in mid-life was positively associated with number of children (Schelbaum et al, 2021). It can be challenging to separate intrinsic biological factors from social/environmental factors concerning the post-partum duration of brain and behavior changes in humans, due to the extended period of child rearing and attendant socioeconomic and cultural stressors. Studies in animals, however, in which lifespans are shorter and environmental conditions can be better controlled, can help to address these factors.

Another relatively neglected aspect of research has been the consequences of reproductive experience in males (Horrell et al, 2019). Although reproductive experience (e.g., mating, rearing of offspring) has less obvious effects on male than female physiology, some evidence in rodents suggests that male reproductive experience can alter prefrontal cortical neural morphology (Wang et al, 2018), hippocampal neurogenesis, and oxytocin signaling in several brain regions, particularly in species in which males engage in care for offspring (Horrell et al, 2021). In humans, brain imaging in fathers during the first 4 months postpartum reveals an increase in grey matter volume in the striatum, amygdala, hypothalamus, and lateral PFC, but a decrease in grey matter volume in the OFC and posterior cingulate cortex (Kim et al, 2014). Studies of male reproductive experience on behavior have largely targeted paternal behavior toward offspring (Abraham & Feldman, 2022; Guoynes & Marler, 2020; Rilling & Mascaro, 2017), response to stressors (Chauke et al, 2011; Harris et al, 2011; Zhao et al, 2017), and paternal aggression (Bukhari et al, 2019; Schneider et al, 2003; Trainor & Marler, 2001). To our knowledge, however, there have been no assessments of the effects of male reproductive experience on executive functions such as cost-benefit decision-making, despite the evidence for physiological consequences of reproduction on the male brain.

The goal of the current studies was to begin to address these gaps in our understanding of the effects of reproductive experience on cognition. To do so, we evaluated the long-term effects of reproductive experience in both female and male rats on several forms of cost-benefit decision making that have been linked previously to differences in risks of psychiatric disorders (Amlung et al, 2019; Gowin et al, 2013; Kaye et al, 2013; Linnet et al, 2011; Mitchell, 2019; Soutschek et al, 2022; Steinglass et al, 2017). On each task, reproductively-experienced and -naïve rats were given choices between a small reward associated with no cost and a large reward associated with various possible costs (delay to delivery, explicit punishment, reward omission). Complementary experiments were conducted to evaluate effects of reproductive experience on food motivation, shock reactivity threshold, and cognitive flexibility.

## 2. Methods

### 2.1 Subjects

Female (n=32) and male (n=24) Long-Evans rats, 50 days of age upon arrival from Charles River Laboratories, were individually housed on a 12 h light/dark cycle (lights on at 0700), maintained at a consistent temperature of 25lJ, and had access to water and 2919 Teklad irradiated global 19% protein chow. Rats were allowed to acclimate to vivarium conditions for at least one week before the start of any procedures.

Experiments were conducted in two cohorts. In the first cohort (females, n=16, males, n=8), half of the females (n=8) were paired with the males while the rest of the females (n=8) remained unpaired. Males were removed from the cages once pregnancies were confirmed. For the rest of this report, the paired (mated) rats are referred to as reproductively experienced (RE), and the unpaired rats as reproductively naïve (RN). RE females gave birth, nursed, and pups were weaned after 21 days. One week after weaning, RE females were paired again with a different male from the same group (to minimize potential effects of specific male mates on females’ reproductive experience). Males were removed again once pregnancies were confirmed, and were not used further in this cohort. Females gave birth, nursed, and pups were weaned after 21 days. One female from this cohort did not become pregnant in either round of pairing and was excluded from the study.

In the second cohort (females, n=16, males, n=16), half of the females (n=8) were paired with half of the males (n=8) while the rest of the females (n=8) and males (n=8) remained unpaired. Males were separated once pregnancies were confirmed. RE females gave birth, nursed, and pups were weaned after 21 days. One week after weaning, RE females were paired again with a different male from the paired male group. Males were removed again once pregnancies were confirmed. Females gave birth, nursed, and pups were weaned after 21 days. One female from this cohort was eliminated from the study due to infanticide in both of her litters. RE and RN males participated in behavioral experiments in the second cohort.

All rats had access to food and water *ad libitum* until one week after the last weaning, when they were put on food restriction. Water access remained *ad libitum* at all times. Behavioral experiments started once rats reached 85% of their free-feeding weights, and rats were individually housed throughout the duration of the behavioral experiments. The behavioral tasks in which the rats were tested proceeded in the following order for both cohorts: intertemporal choice, progressive ratio schedule of reinforcement, risky decision-making, probabilistic reversal learning, reward omission vs. punishment, and shock reactivity threshold, and took approximately six months to complete. All procedures were conducted in accordance with the University of Florida Institutional Animal Care and Use Committee and adhered to the guidelines of the National Institutes of Health.

### 2.2 Apparatus

Behavioral testing was conducted in eight standard operant chambers (Coulbourn Instruments). Chambers were contained in sound-attenuating cubicles, and were computer-controlled through Graphic State 4.0 software (Coulbourn Instruments). Locomotor activity was monitored via infrared motion detectors installed on the ceilings of the chambers. Each operant chamber was equipped with a food trough containing a photobeam sensor to detect nosepokes into the trough, two retractable levers (one on each side of the food trough), a feeder installed on the outside wall of the chamber and connected to the food trough to deliver 45Lmg purified ingredient rodent food pellets (Test Diet; 1811155 5TUM), and a stainless steel floor grate connected to a shock generator that could deliver scrambled footshocks. Each sound-attenuating cubicle included a house light mounted on the rear wall (outside of the operant chamber).

### 2.3 Behavioral procedures

#### 2.3.1 Risky decision-making task

Prior to the start of all operant behavioral testing, rats were trained on a sequence of shaping protocols to learn how to retrieve food from the food trough, nosepoke in the food trough to initiate a trial, and press the levers to obtain food. Shaping procedures are described in detail in (Blaes et al, 2022).

In the risky decision-making task (RDT; (Simon et al, 2009)), rats made discrete-trial choices between two levers. Each trial started with illumination of the house light and the food trough light. A nosepoke into the food trough during this time caused the food trough light to be extinguished and either one lever (on forced choice trials) or both levers (on free choice trials) to extend into the chamber. A failure to nosepoke within 10 s was counted as an omission. A press on one lever (the “small, safe” lever) yielded a single food pellet, while a press on the other lever (the “large, risky” lever) yielded two food pellets and was accompanied by varying probabilities of a mild footshock (1.0 s in duration). A failure to press either lever within 10 s resulted in termination of the trial and marked it as omission. Lever presses were followed by retraction of the lever(s), illumination of the food trough light, and delivery of food pellets. The food trough light was extinguished upon retrieval of the pellets or after 10 s, whichever came first. Trials were separated by an intertrial interval (ITI) in which the house light was extinguished. Each session was comprised of 5 blocks of trials, with different probabilities of shock accompanying the large reward in each block (0, 25, 50, 75, 100%). Figure 1A depicts a schematic of the RDT. Each block of trials began with 8 forced-choice trials (4 for each lever) in which only one lever was extended. The purpose of these trials was to remind rats of the probability of shock in that block. The forced choice trials were followed by 10 free-choice trials in which both levers were extended. Sessions in the RDT were 60 minutes in duration and consisted of 90 trials, each 40 s long. The left/right positions of the small and large reward levers were randomized across rats, but remained consistent for each rat over the course of the task. Shock intensities were 150 µA for the first cohort and 250 µA for the second cohort, but were the same for all rats within each cohort. Rats were trained on the RDT until stable choice performance emerged (see section 2.4, Data Analysis for definition of stable performance).

**Figure 1.**
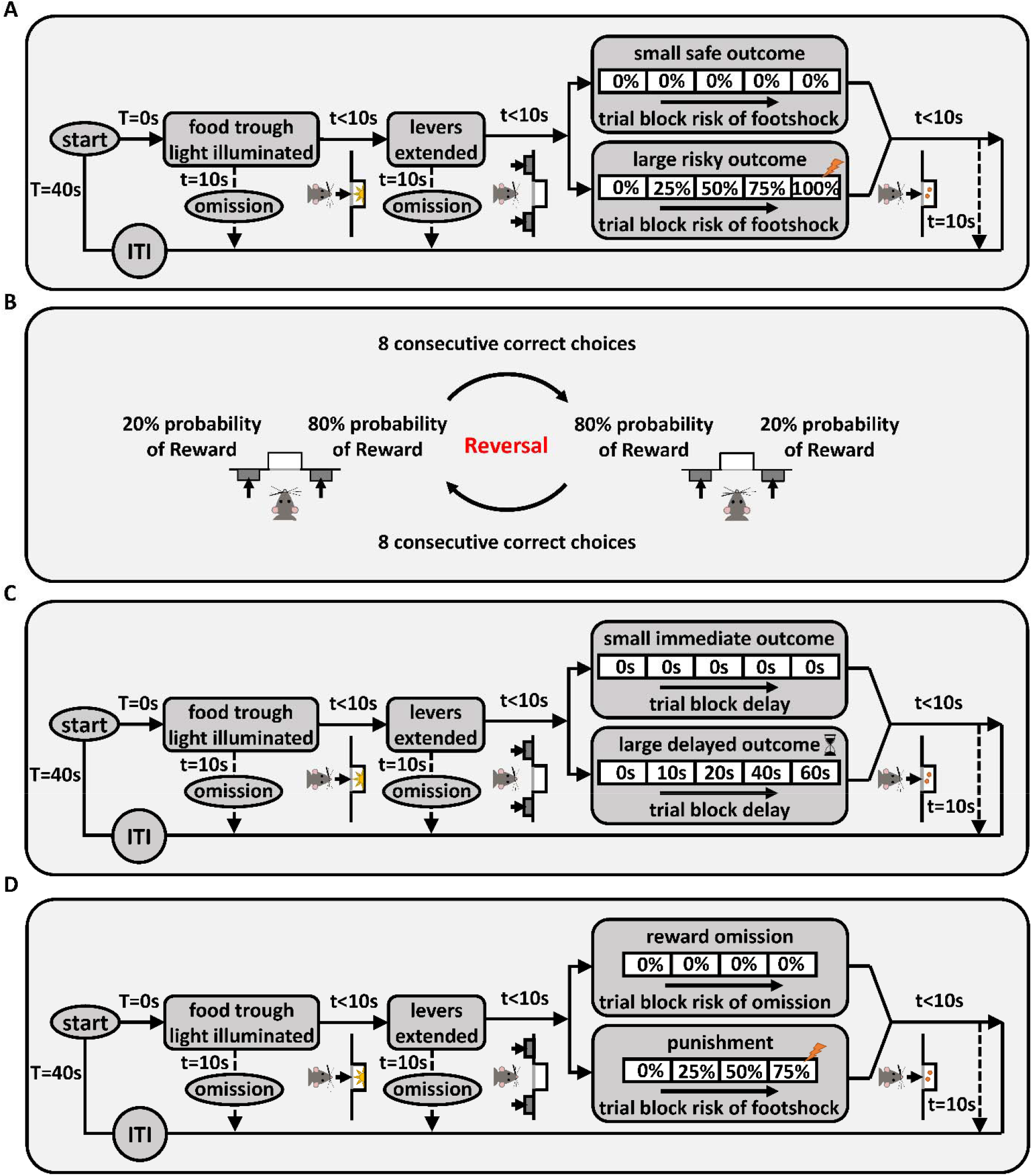
Behavioral task schematics. **(A) Risky decision-making task (RDT).** Each trial was initiated by nosepoking into the food trough, and pressing a lever led to delivery of food pellet(s). If the large reward lever was chosen, delivery of this reward was accompanied by probabilistic shock delivery. There were five blocks of trials, and the probability of shock increased as blocks proceeded. “T” denotes the total time spent in the trial and “t” denotes the time spent in each state of the trial. **(B) Probabilistic Reversal Learning.** The left/right positions of the advantageous (80% chance of reward) and disadvantageous (20% chance of reward) levers were reversed once rats chose the advantageous side 8 times consecutively. **(C) Intertemporal Choice Task**. Each trial was initiated by nosepoking into the food trough, and pressing a lever led to delivery of food pellet(s). If the large reward lever was chosen, food delivery was preceded by a delay. There were five blocks of trials in the task, in which the delay to large reward delivery increased across successive blocks. **(D) Reward Omission vs. Punishment Task (ROVP)** Rats had to decide between risk of reward omission vs. risk of punishment. Each trial was initiated by nosepoking into the food trough, and pressing the levers led to either delivery of reward alone, reward omission, or a reward accompanied by a footshock. There were five blocks of trials in the ROVP task, in which the probability of contingencies (reward omission or shock accompanying the reward) increased across successive blocks.

#### 2.3.2 Progressive Ratio Schedule of Reinforcement

Motivation to pursue food was tested in a progressive ratio schedule of reinforcement task in the same operant chambers used for the RDT. In this task, a single lever (the same lever used as the “small, safe” lever in the RDT) was extended into the chamber and remained extended for the duration of the session. Presses on the lever earned a single food pellet reward; however, the number of presses required to earn subsequent food rewards increased with each successive reward earned according to the following sequence: 1, 3, 6, 10, 15, 20, 25, 32, 40, 50, 62, 77, 95, 118, 145, 175, 205, 235, 265, 295, 325, 355. In any trial, failure to earn a food pellet within 10 minutes resulted in termination of the session. Testing in this task continued for 10 sessions (1 session/day).

#### 2.3.3 Shock Reactivity Threshold

To determine whether reproductive experience affects reactivity to shock, rats were tested in a shock reactivity threshold assay (Bonnet & Peterson, 1975; Orsini et al, 2017a). Rats were placed in a standard operant chamber (a different chamber from that in which they underwent the other behavioral tests) and were initially administered a 400 µA shock to attenuate spontaneous movement in the chamber. Subsequent shocks (1.0 s in duration) were delivered at 10 s intervals, beginning at 50 µA. Responses to the shocks were recorded by a trained observer. Shock reactivity criteria were defined as 1) flinch of a paw or a startle response, 2) elevation of one or two paws, 3) rapid movement of three or all paws. If a shock did not elicit any of the three behaviors, the intensity was increased by 25 µA. If a response was elicited, the intensity was reduced by 25 µA. This procedure continued until the rat was responsive to an intensity 3 times and not responsive to the intensity 25 µA lower twice in a row (following a “yes-no-yes-no-yes” pattern). If a rat showed three consecutive reactions to 50 µA, then 50 µA was recorded as the reactivity threshold for that rat. Each rat was tested twice (on two consecutive days) and the results were averaged.

#### 2.3.4 Probabilistic Reversal Learning

To evaluate effects of reproductive experience on behavioral flexibility, rats were tested on a probabilistic reversal learning task (PRT). In this task (which was modeled on (Dalton et al, 2014) and conducted in the same operant chambers used for the RDT), rats were presented with two levers that differed in the probability with which a press yielded a food reward (Figure 1B). Each 15 s trial began with illumination of the house light for 3 s, followed by extension of both levers. Pressing either lever led to retraction of both levers, illumination of the food trough light, and delivery of a single food pellet (on rewarded trials), followed by an ITI. On trials that were not rewarded (no food delivery), a lever press was immediately followed by the ITI. Failure to press either lever within 10 s of their extension caused the levers to be retracted and the trial counted as an omission. At the start of each session, the probability of food delivery on one of the levers (left or right, randomized across sessions) was 80% (advantageous side) and the probability on the other was 20% (disadvantageous side). Eight consecutive choices of the advantageous side caused the probabilities of reward delivery on the two levers to reverse, such that the previously advantageous side became disadvantageous and vice versa, and rats had to learn the new reward contingencies. This criterion continued throughout the 200 trials in a session, such that a rat could achieve multiple reversals within a session. Each session was 50 min in duration, and rats were tested in the task for 10 consecutive sessions (1 session/day).

#### 2.3.5. Intertemporal Choice Task

Preference for immediate vs. delayed gratification was assessed in an intertemporal choice task (Evenden & Ryan, 1996; Hernandez et al, 2020). In this task (conducted in the same operant chambers used for the RDT), rats made discrete-trial choices between a small, immediate reward and a large, delayed reward. Each trial started with illumination of the house light and the food trough light. A nosepoke into the food trough during this time caused the food trough light to be extinguished and either one lever (on forced choice trials) or both levers (on free choice trials) to extend into the chamber. A failure to nosepoke within 10 s was counted as an omission. Pressing the small reward lever led to illumination of the food trough light and immediate delivery of a food pellet. Pressing the large reward lever led to the delay phase in which, after the delay timer expired, the food trough light was illuminated and three food pellets were delivered. A failure to press a lever within 10 s was counted as an omission. Lever presses were followed by retraction of the lever(s). The food trough light was extinguished upon food retrieval or after 10 s, whichever came first. Trials were separated by an ITI in which the house light was extinguished. Each session was comprised of 5 blocks of trials with different delay durations (0, 10, 20, 40, 60 s). Figure 1C shows a schematic of the task. Each block of trials began with 2 forced-choice trials (one for each lever) in which only one lever was extended. The purpose of these trials was to remind rats of the duration of the delay in that block. The forced-choice trials were followed by 10 free-choice trials in which both levers were extended. Sessions in the intertemporal choice task were 80 minutes in duration and consisted of 60 trials, each 80 s long. The left/right positions of the small and large reward levers were randomized across rats, but remained consistent for each rat over the course of the task. Rats were trained on the intertemporal choice task until stable choice performance emerged (see section 2.4 Data Analysis for definition of stable performance).

#### 2.3.6. Reward Omission vs. Punishment Task

To evaluate preference for different types of risks, rats were tested on a novel task in which they made choices between two parallel contingencies: risk of reward omission vs. risk of punishment (the “Reward Omission Vs. Punishment” (ROVP) task). Each trial in this task started with illumination of the house light and the food trough light. A nosepoke into the food trough during this time caused the food trough light to be extinguished and either one lever (on forced choice trials) or both levers (on free choice trials) to extend into the chamber. A failure to nosepoke within 10 s was counted as an omission. A press on one lever (the “omission” lever) yielded a single food pellet that was delivered with varying probabilities, while a press on the other lever (the “punishment” lever) always yielded a single food pellet but was accompanied by varying probabilities of a mild footshock. A failure to press either lever within 10 s resulted in termination of the trial and was counted as an omission. Lever presses were followed by retraction of the lever(s), illumination of the food trough light, and delivery of a food pellet. The food trough light was extinguished upon retrieval of the pellet or after 10 s, whichever came first. Trials were separated by an ITI in which the house light was extinguished. Each session was comprised of 4 blocks of trials with different probabilities of contingencies (0, 25, 50, and 75% probability of reward omission or shock delivery). Figure 1D shows a schematic of the ROVP task. Each block of trials began with 8 forced-choice trials (4 for each lever) in which only one lever was extended. The purpose of these trials was to remind rats of the probability of contingencies in that block. The forced-choice trials were followed by 10 free-choice trials in which both levers were extended. Sessions in the ROVP task were 48 minutes long and consisted of 72 trials, each 40 s in duration. The left/right positions of the “reward omission” and “punishment” levers were randomized across rats, but remained consistent for each rat over the course of the task. Shock intensities were the same for all rats within each of the two cohorts (200 µA and 350 µA for the first and second cohort respectively, 1.0 s in duration). Rats were trained on the ROVP task until stable choice performance emerged (see section 2.4 Data Analysis for definition of stable performance).

### 2.4. Data analysis

Data were collected and processed using custom protocols and analysis templates in Graphic State 4.0. Statistical analyses were conducted and graphs created using GraphPad Prism 9. To evaluate stable performance in the decision-making tasks, two-factor repeated measures analyses of variance (ANOVA, with session and trial block as within-subjects factors) were conducted on choice data across three (RDT and ROVP) or five (intertemporal choice task) consecutive sessions within each of the four groups (male and female, RE and RN). Stability was defined as the absence of a significant main effect of session or session x trial block interaction.

The primary measures of interest in the decision-making tasks were as follows: In the RDT, percentage of large, risky reward lever presses in each block (out of the total number of trials completed); In the intertemporal choice task, percentage of large, delayed reward lever presses in each block (out of the total number of trials completed); In the ROVP task, percentage of either reward omission lever presses, punishment lever presses, or no lever presses (omitted trials), as a proportion of the total possible number of trials in each block. For rats of each sex, these primary measures were evaluated using a two-factor repeated-measures ANOVA, with reproductive experience as a between-subjects factor and trial block as a within-subjects factor, on data averaged across stable sessions. In the intertemporal choice task, choice indifference points were calculated using a hyperbolic decay function fitted to each rat’s percent choice of the large reward at each delay block. The resulting formula was then used to estimate the delay at which choice of the large reward was equal to 50% (the delay at which the rat was equally likely to choose the large or small reward).

To examine the immediate effects of aversive outcomes on trial-by-trial choice behavior under conditions in which significant differences in choice behavior were observed, win-stay/lose-shift analyses were conducted on data from the RDT and ROVP tasks as in (Orsini et al, 2017a). Trials on which choices were not accompanied by a contingency (footshock on the large reward lever in the RDT and punishment lever in the ROVP, and reward omission on the omission lever in the ROVP) were considered a “win”, and trials on which choices were accompanied by a contingency (footshock or reward omission) were considered a “loss”. Win-stay was defined as when a win trial was followed by choice of the same option on the next trial. Lose-shift was defined as when a lose trial was followed by a switch to the opposite choice (to the safe choice in the RDT and to the other lever in the ROVP) on the next trial. Win-stay performance was measured by dividing the number of win-stay trials by the total number of win trials, and lose-shift performance was measured by dividing the number of lose-shift trials by the total number of lose trials. Both measures were calculated from free-choice trials only. Results were compared between the RE and RN female groups using a two-factor repeated-measures ANOVA, with reproductive experience as a between-subjects factor and trial block as a within-subjects factor on data averaged across stable sessions.

Progressive ratio and probabilistic reversal learning data were evaluated using a two-factor repeated-measures ANOVA, with reproductive experience as a between-subjects factor and session as a within-subjects factor, across the 10 days of task performance. Shock reactivity threshold results were analyzed using a Welch’s t-test, with reproductive experience as the grouping factor.

Ancillary measures in the decision-making tasks, such as locomotor activity, shock reactivity (locomotor activity during the shock delivery period), and omissions during free choice trials were analyzed using a Welch’s t-test, comparing data averaged across trial blocks during the stable sessions of performance. Latencies to press levers during forced- and free-choice trials were used to assess motivation to pursue the outcomes associated with each choice. These data were analyzed using a three-factor repeated-measure ANOVA, with reproductive experience as a between-subjects factor and lever identity and trial block as within-subjects factors, on data averaged across stable sessions. For all analyses, the Greenhouse-Geisser correction was used to account for violations of sphericity in ANOVAs, and p values less than or equal to .05 were considered significant.

## 3. Results

### 3.1. Females

#### 3.1.1. Risky decision-making

Rats were tested on the RDT until stable performance emerged (first cohort, 36 sessions, second cohort, 40 sessions). Analysis of % choice of the large, risky reward using a two-factor repeated measures ANOVA (Group x Shock Probability) revealed a main effect of Shock Probability (F(2.32,63.26)=36.71, p<0.01), such that preference for the large reward declined as probability of shock increased, but no main effect of Group (F(1,28)=1.91, p=0.18). There was, however, a significant interaction between Shock Probability and Group (F(4,109)=3.64, p<0.01), such that RE females chose the large reward more frequently than RN females at higher shock probabilities (Figure 2A).

**Figure 2.**
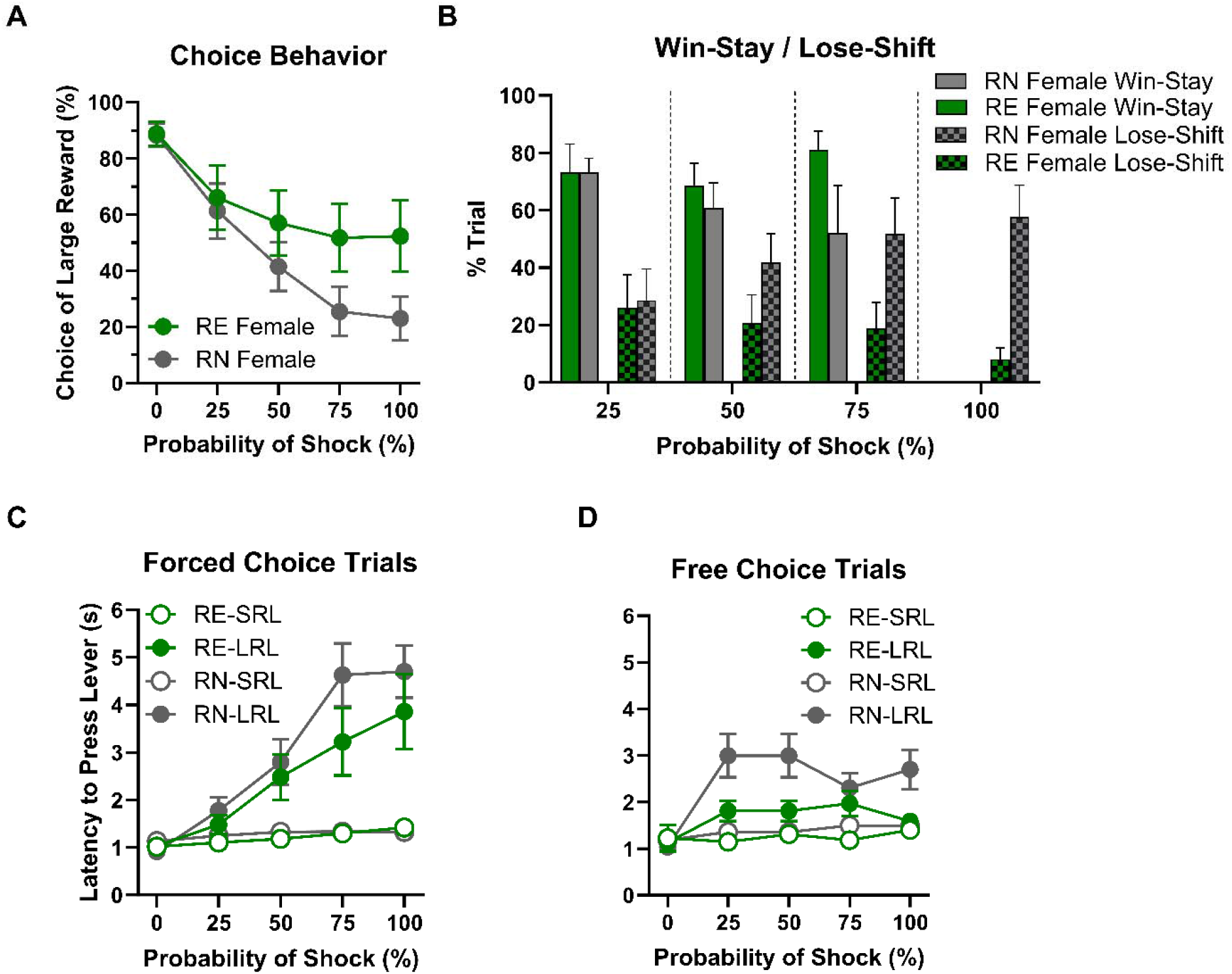
Risky decision making task performance in females. **(A) Choice Behavior**. RE females chose the large risky reward more frequently than RN females at higher shock probabilities. **(B) Win-stay/Lose-shift.** There were no main effects or interaction on win-stay performance. Lose-shift choice, however, was less frequent in RE compared to RN females as shock probabilities increased. **(C)** and **(D) Latencies to press levers.** On forced-choice trials, RE and RN females’ latencies to press the small reward lever (SRL) were comparable, and differences between the group latencies in pressing the large reward lever (LRL) did not reach statistical significance. On free-choice trials, however, RE females showed shorter latencies than RN females at higher shock probabilities. Data are represented as means ± SEM.

Analysis of win-stay results using a two-factor repeated measures ANOVA (Group x Shock Probability; Figure 2B) showed that the likelihood of making the same choice after a “win” was unaffected by Group (F(1,22)=1.52, p=0.23), Shock Probability (F(1.63,27.64)=1.15, p=0.32), or the interaction of the two variables (Group x Shock Probability, F(2,34)=1.34, p=0.28). In contrast, the same analysis conducted on lose-shift results showed that the likelihood of switching to the opposite choice after a loss (shock) was unaffected by Shock Probability (F(2.84,50.11)=2, p=0.13), but there was a main effect of Group (F(1,23)=4.41, p=0.05) and a significant interaction between the two variables (F(3,53)=3.37, p=0.03), such that as shock probability increased across blocks, RN females were more and RE females less likely to switch their choice to the opposite lever following a loss (shock).

Females’ latencies to press levers on both forced-choice and free-choice trials were analyzed using a three-factor repeated-measure ANOVA (Group x Lever x Shock Probability). On forced-choice trials (Figure 2C), latencies were longer at higher shock probabilities (F(2.10,113.5)=46.24, p<0.01) and on the large compared to the small reward lever (F(1, 57)=36.58, p <0.01), particularly at higher shock probabilities (Shock Probability x Lever: F(4,217)=34.18, p<0.01). Response latencies did not differ between RN and RE groups, however (Group: F(1,57)=1.12, p=0.3; Group x Shock Probability: F(4,217)=.86, p=0.49; Group x Lever: F(1,57)=.61, p=0.44; Group x Shock Probability x Lever: F(4,217)=1.59, p=0.18). On free choice trials (Figure 2D), rats had longer latencies at higher shock probabilities (F(1.83,75.43)=20.46, p<0.01) and when choosing the large reward (Lever: F(1,56)=25.71, p<0.01), particularly at higher shock probabilities (Shock Probability x Lever: F(4,165)=12.52, p<0.01). There were no Group x Lever (F(1,56)=2.72, p=0.14) or Group x Lever x Shock Probability (F(4,165)=1.87, p=0.12) interactions; however, there was a main effect of Group (F(1,56)=4.25, p=0.04) and a Group x Shock Probability interaction (F(4,165)=2.79, p=0.03), such that RE females had shorter latencies than RN females, particularly at higher shock probabilities.

A Welch’s t-test was used to evaluate locomotor activity during the ITI, which revealed no difference between the RE and RN females (t(26.49)<0.01, p=0.99). The same analysis conducted on shock reactivity during the task (locomotor activity during shock delivery) and number of free-choice trial omissions also revealed no group differences (shock reactivity: t(18)=0.32, p=0.75; trial omissions: t(27.93)=1.25, p=0.22). Table 1 shows mean (SEM) values across trial blocks between Female groups.

**Table 1.**
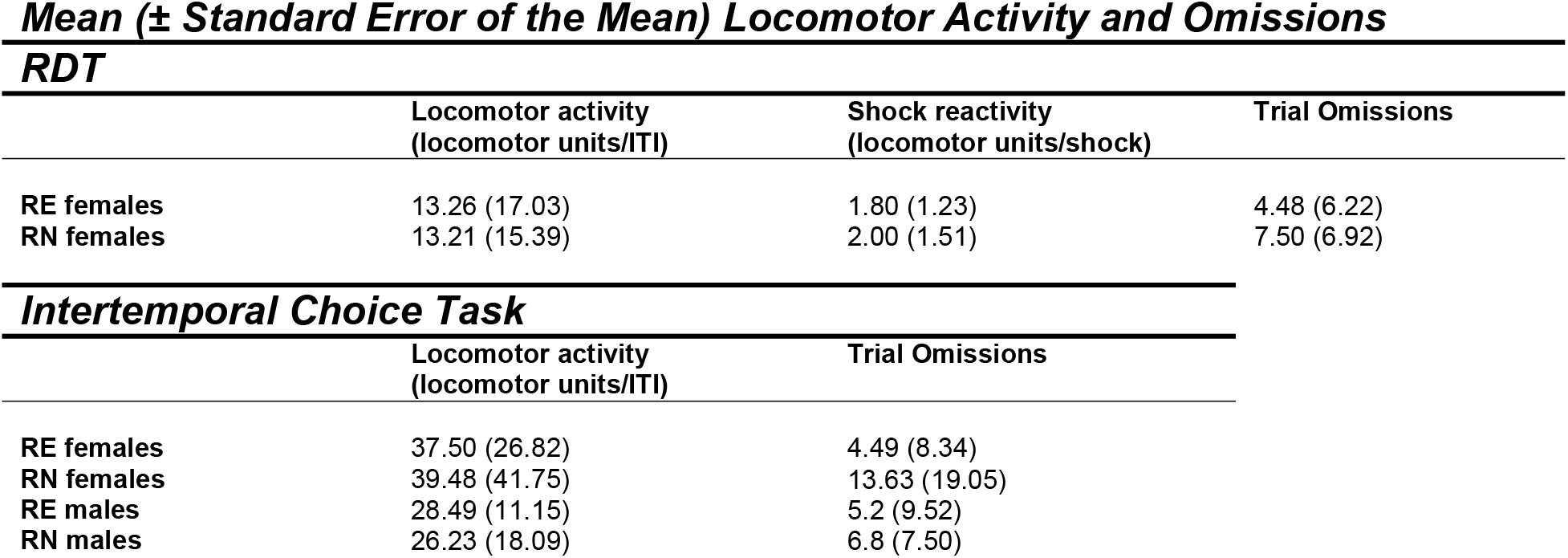
Mean (± Standard Error of the Mean) Locomotor Activity and Omissions RDT.

#### 3.1.2. Progressive Ratio Schedule of Reinforcement

To determine whether the effects of reproductive experience on RDT performance could be due to broader differences in willingness to incur costs to obtain food, rats were tested on a progressive ratio schedule of reinforcement across 10 consecutive sessions (Figure 3A and 3B). A two-factor repeated measures ANOVA (Session x Group) comparing the number of lever presses per session revealed a main effect of Session (F(1.63,45.72)=3.46, p=0.05), but no main effect of Group (F(1,28)=0.48, p=0.49) or interaction between the two variables (F(9,252)=0.71, p=0.70). A comparable analysis conducted on the number of food pellets earned per session revealed similar results (Session: F(1.43,39.96)=2.40, p=0.12; Group: F(1,28)=0.45, p=0.51; Session x Group: F(9,252)=1.17, p=0.32). These data suggest that an increase in food motivation does not account for greater choice of the large, risky reward in RE females in the RDT.

**Figure 3.**
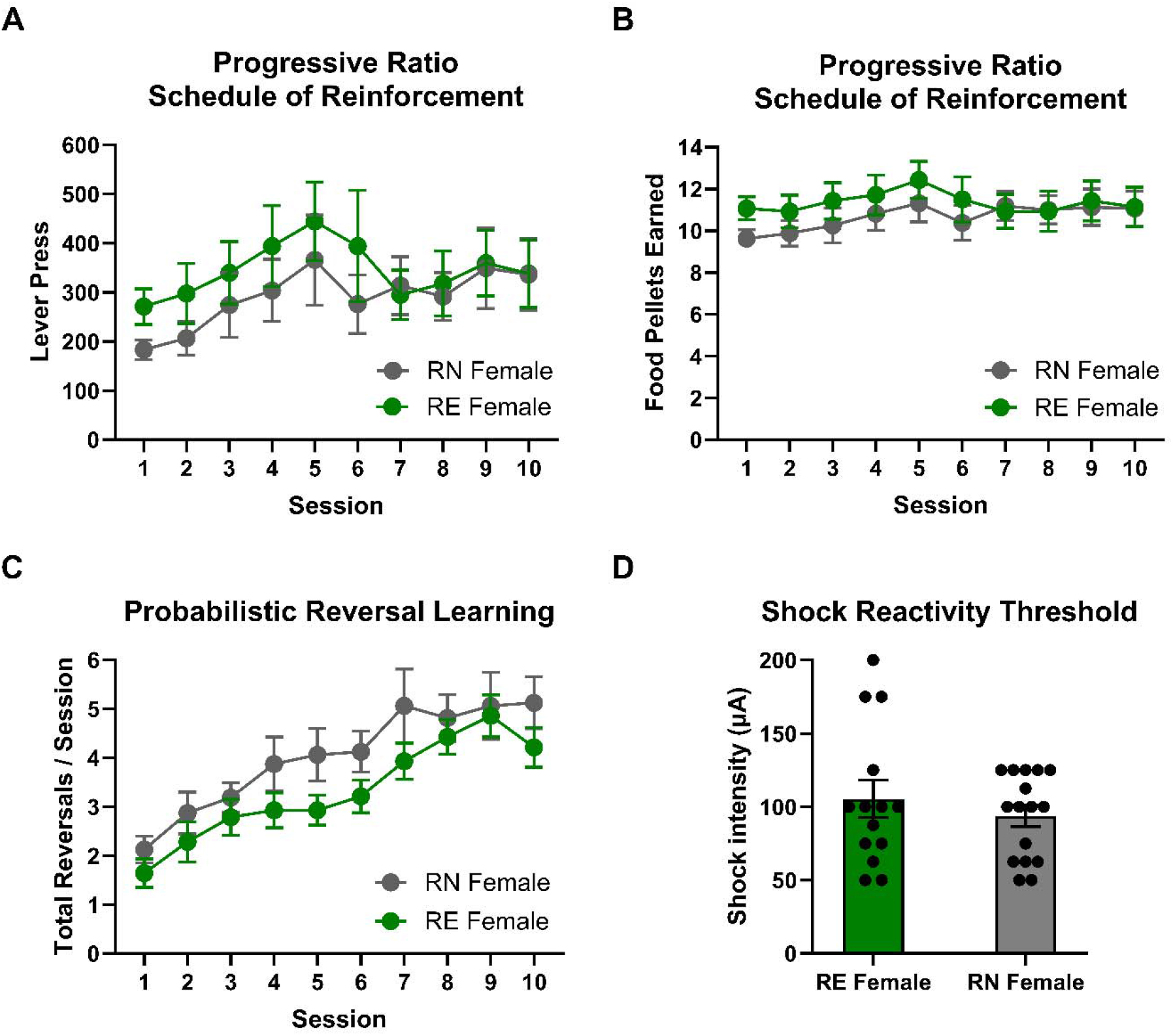
Progressive ratio, reversal learning, and shock reactivity in females. **(A) and (B) Progressive Ratio Schedule of Reinforcement**. There was no group differences in the number of lever presses or food pellets earned between RE and RN females, suggesting equivalent motivation to obtain food. **(C) Probabilistic Reversal Learning**. There were no group differences in the number of reversals completed per session in the probabilistic reversal learning task. **(D) Shock reactivity threshold**. Groups did not differ in shock reactivity threshold. Data are represented as means ± SEM.

#### 3.1.3 Probabilistic Reversal Learning

To determine whether effects of reproductive experience on RDT performance were associated with differences in behavioral flexibility, rats were tested on the probabilistic reversal learning task (Figure 3C). A two-factor repeated measures ANOVA (Group x Session) conducted on the number of completed reversals per session revealed a main effect of Session (F(2.56,71.63)=18.21, p<0.01) such that rats completed more reversals as sessions progressed, but there was neither a main effect of Group (F(1,28)=2.28, p=0.14) nor a Session x Group interaction (F(9,252)=0.50, p=0.87). A similar analysis was conducted on the number of trials completed per reversal (representing the rapidity with which reversals were learned), showing that rats met the reversal criterion in fewer trials as sessions progressed (Session: F(4.03,112.7)=12.87, pL<0.01), but revealing comparable results between groups (Group: F(1,28)=1.82, p=0.19; Session x Group: F(9,252)=0.94, p=0.49). Analysis of the number of total omitted trials per session revealed neither main effects of Group (F(1,28)=0.60, p=0.44) or Session (F(1.94,54.28)=2.57, p=0.09), nor a Session x Group interaction (F(9,252)=1.10, p=0.37). Considered together, these data show that reproductive experience in females does not affect reversal learning, and suggest that reduced behavioral flexibility does not account for greater choice of the large, risky reward in RE rats across blocks in the RDT.

#### 3.1.4 Shock Reactivity Threshold

Rats were tested for their shock reactivity threshold to determine whether the effects of reproductive experience on RDT performance could be due to differences in shock sensitivity (Figure 3D). Comparison of reactivity thresholds using a Welch’s t-test revealed no difference between RE and RN groups (t(20.88)=0.80, p=0.44).

#### 3.1.5. Intertemporal Choice Task

Rats were tested on the intertemporal choice task until stable choice behavior emerged (first cohort, 31 sessions; second cohort, 21 sessions). Comparison of choice of the large, delayed reward in RN and RE rats during stable performance using a two-factor ANOVA (Delay x Group) revealed a main effect of Delay (F(4,112)=144.5, p<0.01) such that rats chose the large reward less frequently as the delay to reward delivery increased, as well as a main effect of Group (F(1,28)=6.59, p=0.02) and a delay x group interaction (F(4,112)=3.52, p<0.01), such that RE rats showed reduced preference for the large reward, particularly at short delays (Figure 4A). There was a notable gap between the RN and RE females’ choice of the large reward in the first block (0 s delay), which was driven by a cluster of RE females (n=5 out of 14) avoiding the large reward even in the absence of delay to its delivery (less than 80% choice of the large reward at the 0 s delay). Removing these rats (as well as one similar outlier from the RN group) closed the gap at the 0 s delay but preserved the significant delay x group interaction (F(4,88)=2.5, p=0.05), with RE females still choosing the large delayed reward less frequently than RN females, most notably at the 10 and 20 s delays. This result might indicate greater sensitivity of RE females to the delay cost associated with the large reward, as the indifference point (50% choice of large delayed reward) for this subgroup of RE females (individuals with more than 80% choice of the large reward at the 0 s delay) was at 10.5 seconds versus 21.4 seconds for the RN group (Figure 4B).

**Figure 4.**
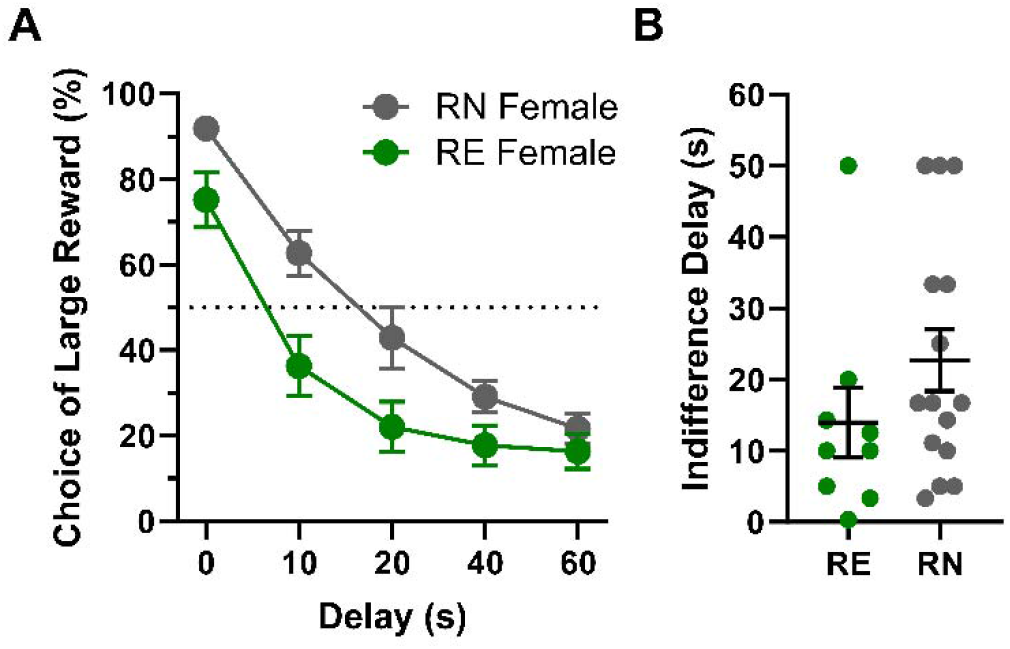
Intertemporal choice task performance in females. **(A) Choice behavior.** RE females showed reduced preference for the large, delayed reward compared to RN females, particularly at short delays. **(B) Indifference delay.** RE females showed shorter indifference delays (the theoretical delay to large reward delivery at which preference between the large, delayed and small, immediate reward is equivalent) compared to RN females. Data are represented as means ± SEM.

Latencies to lever press were analyzed using a three-factor repeated-measure ANOVA (Group x Delay x Lever). On forced-choice trials, there were main effects of Delay (F(2.85,159.8)=21.2, p<0.01) and Lever (F(1, 56)=10.94, p <0.01), as well as a Delay x Lever interaction (Delay x Lever: F(4,224)=29.43, p<0.01), such that latencies were longer on both the large reward lever and at longer delays, and the difference in latencies between the two levers was larger at longer delays. There were no main effects or interactions involving Group, however, (Group: F(1,56)=0.11, p=0.74; Group x Delay: F(4,224)=0.81, p=0.52; Group x Lever: F(1,56)=0.19, p=0.67; Group x Lever x Delay: F(4,224)=1.19, p=0.32). Analyses of latencies on free-choice trials revealed results similar to those on forced-choice trials, with main effects of Delay (F(2.55,139.6)=55.01, p<0.01) and Lever (F(1,56)=17.19, p<0.01) and a Delay x Lever interaction (Delay x Lever: F(4,219)=15.75, p<0.01), but no main effects or interactions involving Group (Group: F(1,56)=0.34, p=0.56; Group x Delay: F(4,219)=0.78, p=0.54; Group x Lever: F(1,56)=0.15, p=0.7; Group x Lever x Delay: F(4,219)=0.39, p=0.81).

A Welch’s t-test comparing locomotor activity in the intertemporal choice task showed no significant difference in activity between RE and RN females (t(25.85)=0.16, p=0.88). The same analysis conducted on the number of omitted trials showed that RE females omitted fewer trials than RN females (t(21.55)=2.50, p=0.02). Table 1 shows the mean (SEM) values for locomotor activity and omitted trials between female groups.

#### 3.1.6. Reward Omission vs. Punishment Task

To determine the effects of reproductive experience on decisions between two different types of risk, rats were tested in a novel behavioral task in which they chose between rewards of the same magnitude (one food pellet) that were associated with different costs (risk of reward omission vs. risk of punishment; the ROVP task).

Females were tested on the ROVP until stable behavior emerged (first cohort, 7 sessions; second cohort, 24 sessions; Figure 5). All rats chose the punished option less frequently as the probability of the contingencies increased across blocks of trials (F(1.54,43.25)=18.7, p<0.01). Choice of the punished option was numerically greater in RE compared to RN females, but this difference did not reach statistical significance (Group: (F(1,28)=3.99, p=0.06; Group x Probability: (F(3,84)=2.27, p=0.09). Choice of the reward omission option also decreased across blocks of trials (F(1.88,52.75)=12.33, p<0.01), but, as with choice of the punished option, it did not differ between groups (Group: F(1,28)=2.79, p=0.11; Group x Probability: F(3,84)=0.82, p=0.49). Finally, trial omissions increased substantially across blocks of trials (F(1.94,54.28)=36.50, p<0.01), but did not differ between groups (Group: F(1,28)=0.71, p=0.41; Group x Probability: F(3,84)=0.92, p=0.43).

**Figure 5.**
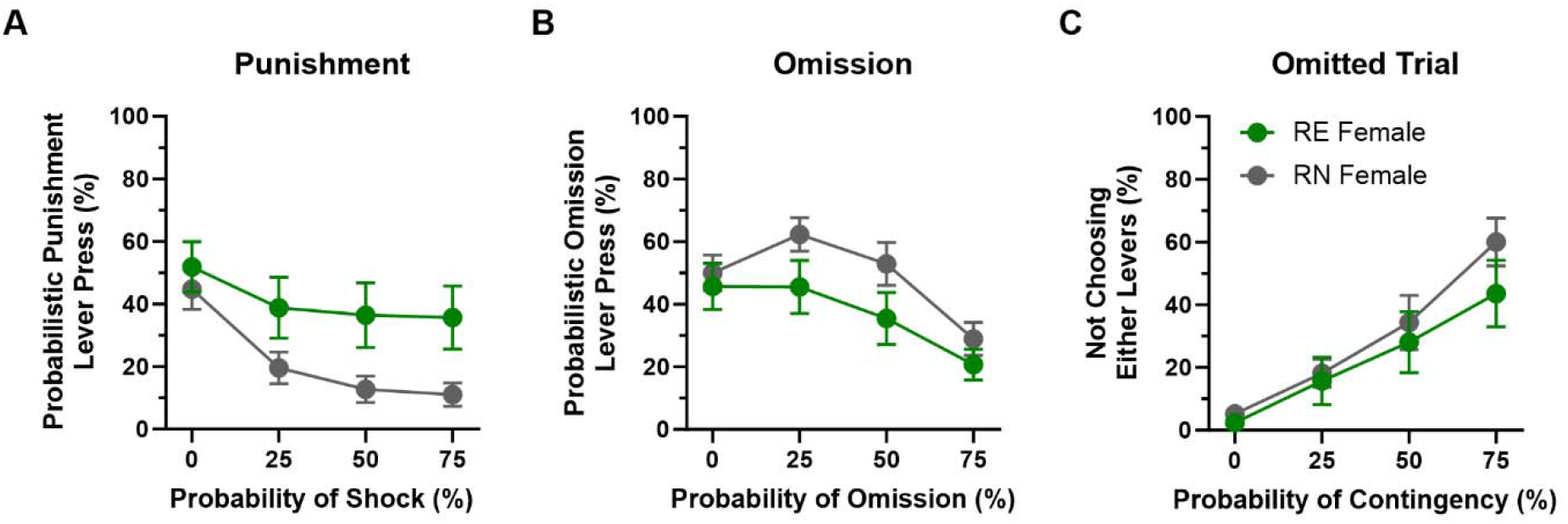
Reward Omission vs. Punishment task performance in females. **(A)** RE females chose the punishment option more frequently than RN females, but this difference was not statistically reliable. **(B)** There were no significant group differences in preference for the reward omission option. **(C)** The number of omitted trials increased across blocks, but was comparable between the two groups. Data are represented as means ± SEM.

Analyses of win-stay/lose-shift behavior in the ROVP task (two-factor repeated measures ANOVA, Group x Probability) revealed a main effect of reproductive experience on win-stay performance on the punishment side (Group, F(1,20)=4.95, p=0.04), such that RE females repeated their choice of the punishment option after a win trial more frequently than RN females. No other significant main effects or interactions were observed with either the punishment or the reward omission option (Table 2).

**Table 2.** Win-Stay/Lose-Shift performance in the ROVP (Females)

|  | **Punishment** |  | **Reward Omission** |  |
| --- | --- | --- | --- | --- |
|  | **Win-Stay** | **Lose-shift** | **Win-Stay** | **Lose-shift** |
| **Reproductive Experience** | F (1, 20) = 4.95,<br>p=0.04 | F (1, 20) = 1.73,<br>p=0.20 | F (1, 28) =0.21,<br>p=0.65 | F (1, 28) = 1.18,<br>p=0.29 |
| **Probability of Contingency** | F (1.83, 21.06) =<br>0.94,<br>p=0.92 | F (1.71, 25.63) =<br>1.55,<br>p=0.23 | F (1.99, 42.8) = 1,<br>p=0.97 | F (1.87, 46.66) =<br>1.12,<br>p=0.33 |
| **Reproductive Experience X Probability of Contingency** | F (2, 23) =0.45,<br>p=0.65 | F (2, 30) = 0.11,<br>p=0.89 | F (2, 43) =0.74,<br>p=0.48 | F (2, 50) =0.88,<br>p=0.42 |

### 3.2. Males

#### 3.2.1 Risky Decision-making Task

To determine the effects of reproductive experience on risk-taking behavior in males, RN and RE males were tested in the RDT following attainment of stable performance (48 sessions). As shown in Figure 6A, a two-factor repeated measures ANOVA (Shock Probability x Group) revealed the expected main effect of Shock Probability (F(1.58,21.66)=10.31, p<0.01) but neither a main effect of Group (F(1,14)<0.01, p=0.99) nor a Shock Probability x Group interaction (F(4,55)=0.38, p=0.82), suggesting that reproductive experience in male rats does not alter decision making involving risk of explicit punishment.

**Figure 6.**
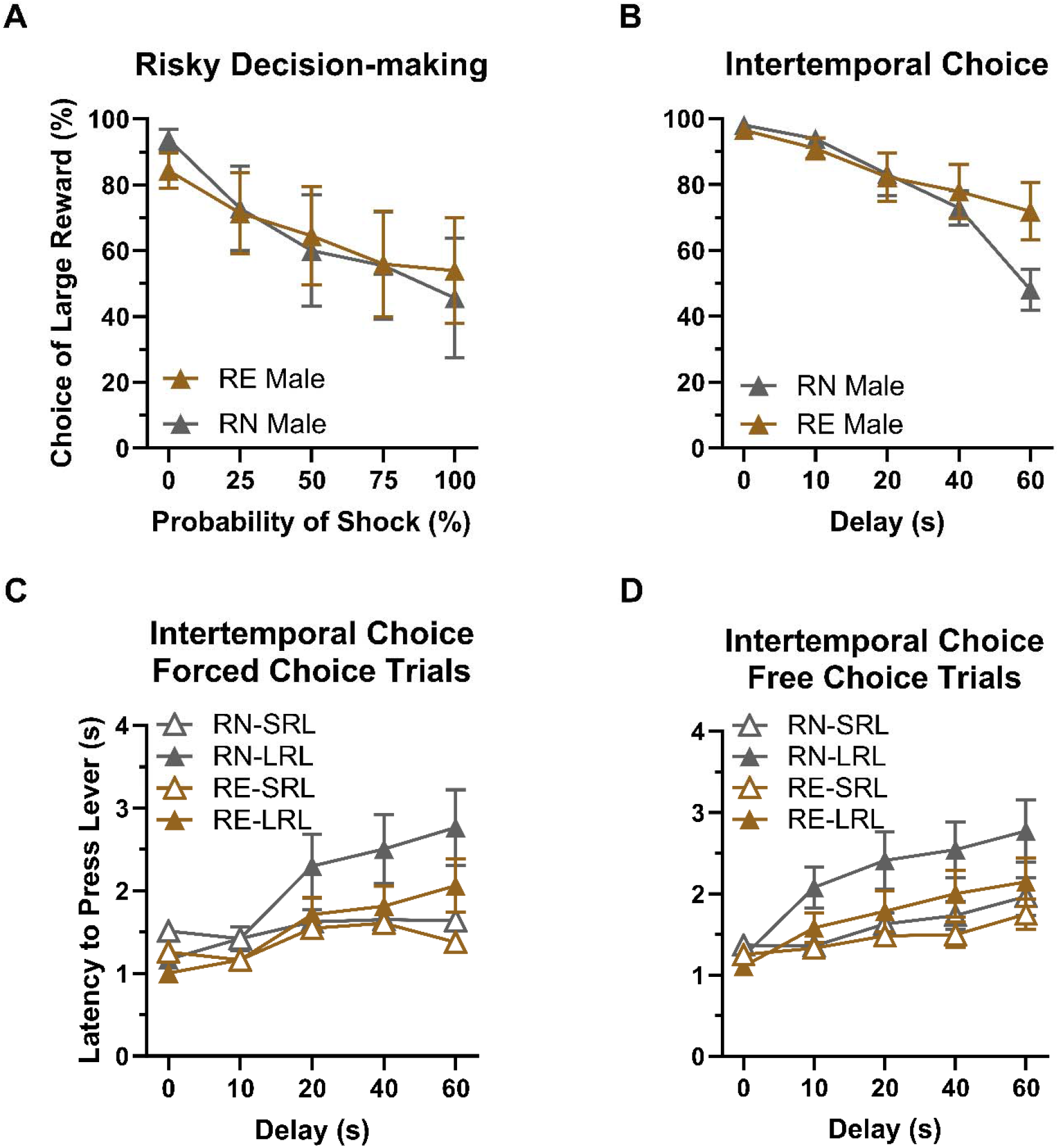
Risky decision making and intertemporal choice task performance in males. **(A) Risky Decision-making in males.** RE and RN males did not differ in risk-taking behavior. **(B) Intertemporal choice task.** RE males showed greater preference for the large, delayed reward compared to RN males, particularly at long delays. **(C) and (D) Latency to press levers in the intertemporal choice task.** There were no significant group differences in RE and RN males’ latencies to press the small reward lever (SRL) or large reward lever (LRL), on either forced- or free-choice trials. Data are represented as mean % choice ± SEM.

#### 3.2.3 Intertemporal Choice Task

Rats were tested on the intertemporal choice task until stable choice behavior emerged (23 sessions, Figure 6B). Comparison of choice of the large, delayed reward in RN and RE rats during stable performance using a two-factor repeated measures ANOVA (Delay x Group) revealed a main effect of Delay (F(1.94,27.13)=28.41, p<0.01) such that rats chose the large reward less frequently as the delay to reward delivery increased. There was no main effect of Group (F(1,14)=0.53, p=0.48), but a Delay x Group interaction (F(4,56)=4.04, p<0.01) was observed, such that RE rats showed greater preference for the large reward specifically at the longest delay.

To further explore the difference in behavior on the intertemporal choice task, males’ latencies to lever press were analyzed using a three-factor repeated-measure ANOVA (Group x Delay x Lever). On forced-choice trials (Figure 6C), there was the expected main effect of delay (F(4,112)=23.93, p<0.01) as well as a Delay x Lever interaction (Delay x Lever: F(4,112)=11.12, p<0.01), such that latencies on the large reward lever increased to a greater extent across delays than latencies on the small reward lever. In contrast, there was no significant main effect of lever (F(1,28)=3.36, p=0.08), nor were there significant main effects or interactions involving Group (Group: F(1,28)=3.76, p=0.06; Group x Delay: F(4,112)=0.55, p=0.70; Group x Lever: F(1,28)=0.79, p=0.38; Group x Lever x Delay: F(4,112)=1.34, p=0.26).

On free-choice trials (Figure 6D), there were significant main effects of delay (F(1.31,36.39)=40.16, p<0.01) and lever (F(1,28)=4.88, p=0.04), as well as a Delay x Lever interaction (Delay x Lever: F(4,111)=8.43, p<0.01) such that latencies increased to a greater extent on the large than the small reward lever across delays. As with the forced-choice trials, however, there were no main effects or interactions involving Group (Group: F(1,28)=2.63, p=0.12; Group x Delay: F(4,111)=1.25, p=0.29; Group x Lever: F(1,28)=0.74, p=0.40; Group x Lever x Delay: F(4,111)=0.76, p=0.56).

A Welch’s t-test was used to evaluate locomotor activity in the intertemporal choice task, and revealed no difference in activity between the RE and RN males (t(11.65)=0.30, p=0.77). The same analysis conducted on the number of omitted trials showed no difference between the RE and RN males (t(13.27)=0.37, p=0.71). Table 1 shows the mean values for locomotor activity and omitted trials across trial blocks between male groups.

#### 3.2.4 Reward Omission vs. Punishment Task

Male RN and RE rats (Figure 7) chose the punished option less frequently as the probability of the contingencies increased across blocks of trials (F(2.2,30.79)=13.04, p<0.01), but there was no difference between the two groups (Group: F(1,14)=0.32, p=0.58; Group x Probability: (F(3,42)=0.89, p=0.46). Choice of the reward omission option also decreased across blocks of trials (F(2.2,30.86)=11.65, p<0.01), but, as with choice of the punished option, it did not differ between groups (Group: F(1,14)=1.21, p=0.29; Group x Probability: (F(3,42)=0.82, p=0.49). Finally, trial omissions increased substantially across blocks of trials (F(1.98,25.79)=35.38, p<0.01), but did not differ between groups (Group: F(1,13)=0.03, p=0.86; Group x Probability: F(3,39)=0.03, p=0.99).

**Figure 7.**
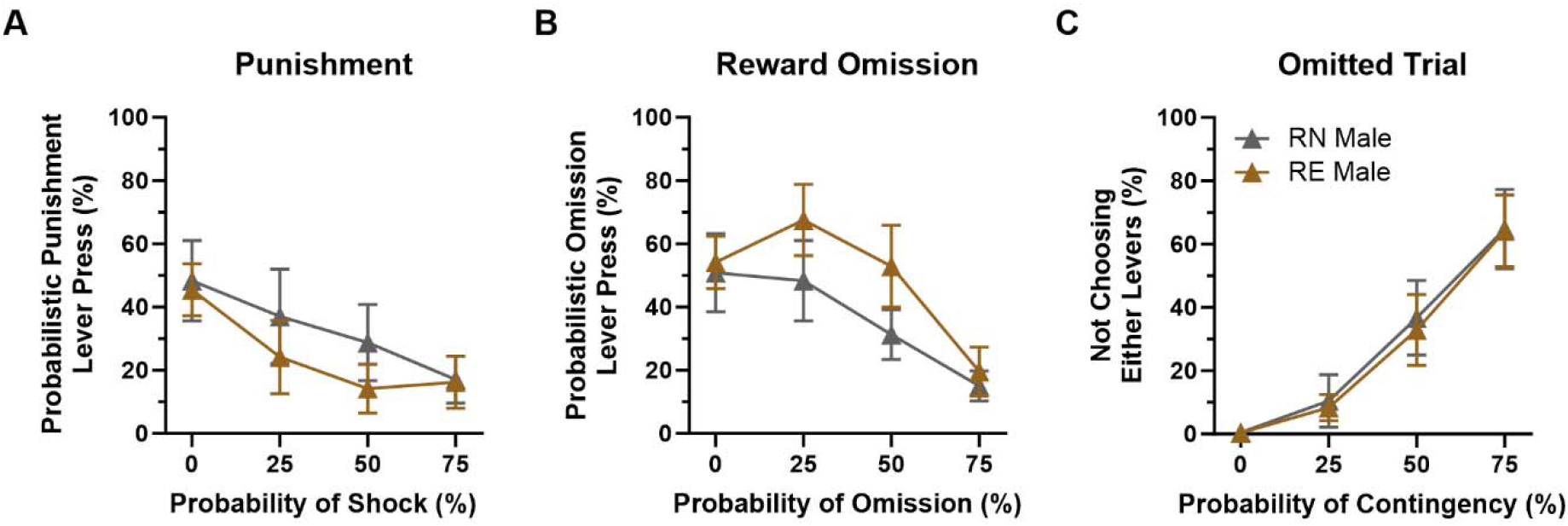
Reward Omission vs. Punishment task performance in males. **(A)** There were no significant group differences in RE and RN males’ choice of the punishment option. **(B)** There were no significant group differences in preference for the reward omission option. **(C)** The number of omitted trials increased across blocks, but was comparable between the two groups. Data are represented as means ± SEM.

#### 3.2.5 Progressive Ratio Schedule of Reinforcement

To determine the effect of reproductive experience on motivation to work for food, male rats were tested on a progressive ratio schedule of reinforcement across 10 consecutive sessions (Figures 8A and 8B). A two-factor repeated measures ANOVA (Session x Group) comparing the number of lever presses revealed no main effects of Session (F(1.71,23.9)=2.44, p=0.12) or Group (F (1,14)=0.96, p=0.34), nor was there an interaction between the two variables (F(9,126)=1.66, p=0.10). A similar analysis of the number of food pellets earned revealed no main effects of Group (F(1,14)=0.74, p=0.41) or Session (F(1.95,27.36)=2.7, p=0.08), but in this case the Group x Session interaction reached statistical significance (F(9,13)=2.25, p=0.02), such that RE males earned more food pellets than RN males in earlier but not later sessions. These data show that reproductive experience in males appears to produce transient increases in food motivation, which might account for the greater preference for the large reward in RE males at the longest delay in the intertemporal choice task.

**Figure 8.**
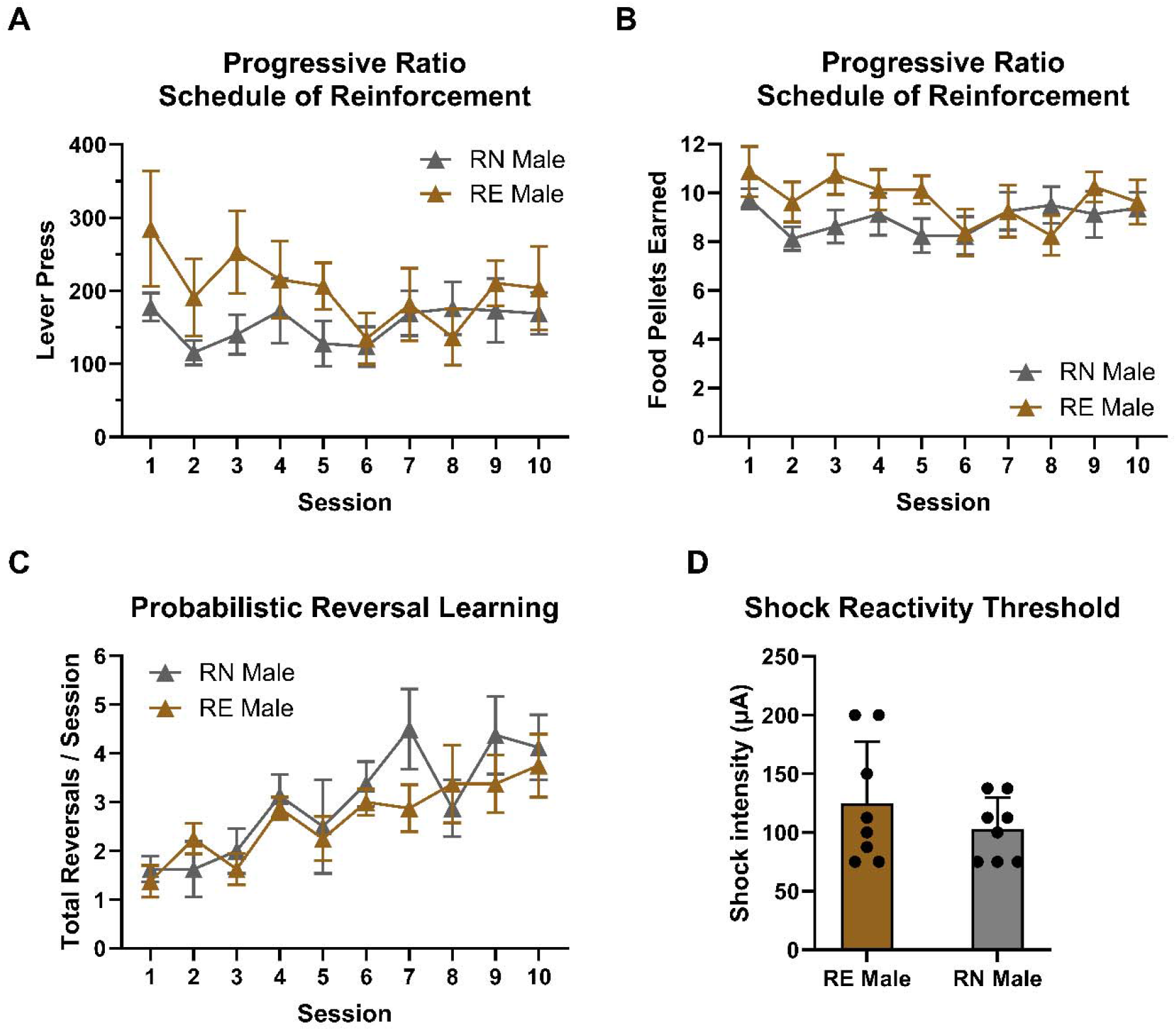
Progressive ratio, reversal learning, and shock reactivity in males. **(A) and (B) Progressive Ratio Schedule of Reinforcement**. Although there was no difference between RE and RN males in the number of lever presses, the interaction between group and session was significant for food pellets earned, such that RE males earned more food pellets than RN males in earlier sessions. **(C) Probabilistic Reversal Learning**. There were no group differences in the number of reversals completed per session in the probabilistic reversal learning task. **(D) Shock reactivity threshold**. Groups did not differ in shock reactivity threshold. Data are represented as means ± SEM.

#### 3.2.7 Probabilistic Reversal Learning

To determine whether reproductive experience affects behavioral flexibility, male rats were tested on the probabilistic reversal learning task (Figure 8C). A two-factor repeated measures ANOVA (Group x Session) conducted on the number of completed reversals per session revealed a main effect of Session (F(3.18,44.46)=7.26, p<0.01) such that rats completed more reversals as sessions progressed, but neither a main effect of Group (F(1,14)=0.47, p=0.50) nor a Session x Group interaction (F(9,126)=0.96, p=0.48). A similar analysis conducted on the number of trials completed per reversal (representing the rapidity with which reversals were learned) showed rats meeting the reversal criterion in fewer trials as sessions progressed (Session: F(3.76,52.65)=6.33, pL<0.01), but no differences between groups (Group: F(1,14)=0.08, p=0.78; Session x Group: F(9,126)=1.52, p=0.15). Analysis of the number of total omitted trials per session revealed no main effect of Group (F(1,14)=0.51, p=0.49), Session (F(2,27.95)=1.38, p=0.27), or a Session x Group interaction (F(9,126)=0.56, p=0.82). Considered together, these data show that reproductive experience in males does not affect reversal learning, and suggest that reduced behavioral flexibility does not account for greater choice of the large, delayed reward in the intertemporal choice task in RE rats.

#### 3.2.6 Shock Reactivity Threshold

Males were tested for their shock reactivity threshold following the same procedures described above for females. A Welch’s t-test revealed no difference in reactivity threshold between RN and RE males (t(10.39)=1.06, p=0.31; Figure 8D).

## 4. Discussion

Plasticity of the female brain during pregnancy and postpartum is widely recognized, yet literature on long term effects of reproductive experience on executive functions is scarce, particularly in animal models in which environmental variables can be tightly controlled, and even more particularly in male subjects. Here we show that the full spectrum of reproductive experience (mating, pregnancy, parturition, and pup rearing) is associated with performance differences across multiple cost-benefit decision making tasks in both females and males. RE females showed greater preference for large, but probabilistically punished reward, and reduced preference for large, delayed over small, immediate reward. In contrast, RE males showed greater preference for large, delayed over small, immediate reward at long delays to reward delivery.

### 4.1 Risky decision making in females

In the RDT, RE females showed greater preference than RN females for the large reward accompanied by probabilistic punishment (i.e., greater risk taking). This difference was not likely due to diminished sensitivity to footshock, as shock reactivity was equivalent in the two groups. There were further no differences between groups in the progressive ratio task, nor were there differences in probabilistic reversal learning, suggesting that differences in food motivation or behavioral flexibility also did not account for the greater risk taking in the RE group. These conclusions are supported by the results from the intertemporal choice task, in which RE females showed reduced preference for large rewards, as well as a more rapid shift toward preference for the small, immediate reward across trial blocks (see Intertemporal Choice section below for further discussion).

Risky decision making in general, and performance in the RDT specifically, are strongly regulated by dopamine signaling (França & Pompeia, 2023; Orsini et al, 2015; Piantadosi et al, 2021). Previous work with the RDT shows that higher levels of risk taking are associated with both higher levels of evoked dopamine release in nucleus accumbens (Freels et al, 2020) and lower expression of D2 receptor mRNA in striatum (Mitchell et al, 2014). Dopamine signaling is also implicated in some aspects of reproduction-related behavior, including food cravings during pregnancy, maternal behavior, and post-partum depression (Haddad-Tóvolli et al, 2022; Rincón-Cortés & Grace, 2020; 2022; Robinson et al, 2011). Recordings of dopamine availability from nucleus accumbens in rats using fast-scan cyclic voltammetry show dopamine transients in mothers when interacting with pups, and that evoked dopamine release under anesthesia is greater in post-partum compared to reproductively naïve rats (Robinson et al, 2011; Shnitko et al, 2017). These findings are consistent with data showing greater sensitivity to dopamine agonists in post-partum compared to reproductively-naïve rats (Byrnes et al, 2001; Byrnes et al, 2011). In humans, PET imaging data show reduced D2/3 receptor availability in striatum post-partum (Moses-Kolko et al, 2012). Given the similarities in dopamine signaling between rats with higher levels of risk taking behavior and post-partum individuals, it seems likely that reproduction-induced shifts in dopamine signaling play a causal role in the elevated risk taking observed in RE females.

In addition to changes in dopamine signaling, the post-partum period is associated with changes in circulating levels of ovarian hormones, including reductions in estradiol and prolactin (Bridges & Byrnes, 2006). Interestingly, ovariectomy (which reduces estradiol levels) causes an increase in risk taking in the RDT, which is partially reversed by estradiol replacement (Orsini et al, 2021). Although the mechanisms by which these manipulations of ovarian hormones cause alterations in risk taking are unclear, it is likely that it occurs at least in part via shifts in dopamine signaling.

### 4.2 Intertemporal choice in females

In the intertemporal choice task, RE females showed reduced preference for the large reward relative to RN females, particularly at shorter (including zero) delays to delivery. Failures to reliably choose the large reward in the absence of delays in intertemporal choice tasks can be interpreted as a deficit in sensitivity to reward magnitude or reduced motivation to obtain the large reward. Inspection of individual rats’ performance at the 0 s delay, however, revealed that when rats from both RE and RN groups that chose the large reward on fewer than 80% of trials at the 0 s delay were excluded, there was equivalent choice of the large reward between the two groups in this block, but the significant interaction between group and delay remained. This (admittedly post-hoc) analysis suggests that reduced preference for the large, delayed reward in RE females is not solely due to reduced sensitivity to the large reward more generally, but rather reflects greater preference for small, immediate over large, delayed reward (i.e., greater impulsive choice). Consistent with this interpretation, RE females omitted fewer trials than RN females in this task and showed intact preference for the large reward in the RDT, not to mention the absence of a difference between the two groups in the progressive ratio task.

Performance on the intertemporal choice task and the RDT is not correlated in rats (Olshavsky et al, 2014; Orsini & Setlow, 2017; Shimp et al, 2015; Simon et al, 2009), but elevated levels of both risk taking and preference for immediate over delayed gratification are associated in some clinical conditions (e.g., substance use disorders (Bornovalova et al, 2005; Costanza et al, 2021; Smith & Cyders, 2016)). The fact that RE females showed this same pattern of performance in the two tasks relative to RN females could suggest that a common mechanism underlies both changes. Postpartum shifts in dopamine signaling are one possible candidate for such changes. Lower levels of striatal D2/D3 receptor availability are associated with greater discounting of delayed rewards in several clinical populations (Ballard et al, 2015; Heinz et al, 2004; Joutsa et al, 2015) and similar findings are evident in untreated rats (Dalley et al, 2007; Joutsa et al, 2015). Given that striatal D2/D3 receptor availability is reduced in postpartum women (Moses-Kolko et al, 2012), such changes in dopamine signaling could account for the shifts in both risky and intertemporal choice observed in the present study.

### 4.3 Cost-benefit decision making in RE males

In contrast to female rats, male rats exhibited no effects of reproductive experience on performance in the RDT. On the intertemporal choice task, however, RE males showed greater choice of the large reward specifically at the longest delay to reward delivery. This pattern could be interpreted as attenuated behavioral flexibility (i.e., an inability to shift choices away from the large reward as its cost increases) or increased reward motivation. The absence of differences between RN and RE males on probabilistic reversal learning argues against attenuated behavioral flexibility; however, the increased responding in the progressive ratio task might indicate greater reward motivation in RE males.

Reproductive experience in male rodents leads to paternal behavior (feeding, huddling, retrieving) and territorial aggression (Bukhari et al, 2019; Saltzman et al, 2017; Schneider et al, 2003; Trainor & Marler, 2001). While these effects are acute consequences of sexual experience and fatherhood, few studies have addressed long-term neurobiological changes in males following sexual behavior (Pitchers et al, 2010; Woller et al, 2012), and to our knowledge, no study has addressed the long-term neurobiological effects of sexual experience on male decision-making.

Sexual experience in male rats acutely enhances dopamine release in nucleus accumbens (Damsma et al, 1992), and can result in an enhanced (sensitized) response to amphetamine as revealed by greater amphetamine conditioned place preference (CPP) in sexually-experienced vs. –naïve males (Pitchers et al, 2010) (see also (Fiorino & Phillips, 1999a; Fiorino & Phillips, 1999b) for related results). Moreover, VTA dopamine neuron activity during sexual experience in male rats is necessary for the sensitizing effects of such experience on subsequent amphetamine CPP (Beloate et al, 2016). If a similar sexual experience-induced sensitized dopaminergic response to rewarding stimuli were in effect in the present study, it might account for the increased preference for the large, delayed reward in RE rats, and would be consistent with their greater number of food pellets earned early in the progressive ratio task. Arguing against this possibility is the fact that no effects of male reproductive experience were evident in the RDT; however, given that the intertemporal choice and progressive ratio tasks were more proximal in time to sexual experience, a potential reward sensitization effect in males could have dissipated by the time of RDT testing.

### 4.4 Reproductive experience and aging

Animal models of aging, similar to other fields of animal research, use reproductively-naive subjects as standard practice, but the extent to which this represents the human condition (in which some form of reproductive experience is nearly ubiquitous) remains to be addressed. Reproductive history has been recognized as a contributing factor to cognitive aging in humans (Harville et al, 2020; Jung et al, 2020; Orchard et al, 2023). For example, a study in humans shows that number of childbirths is a predictor of brain age, with parous females exhibiting “younger-looking” brains in middle age (de Lange et al, 2019). Using MRI data, the same investigators found that more childbirths were associated with less apparent brain aging based on structural characteristic (cortical thickness, area, and volume) in striatal and limbic regions and, in particular, the nucleus accumbens (de Lange et al, 2020). Longer reproductive span and greater number of children are also associated with larger grey volume matter in brain regions vulnerable to Alzheimer disease and cognitive aging in midlife (Schelbaum et al, 2021). Such findings emphasize the importance of including reproductive experience in aging studies.

### 4.5 Limitations and future work

One limitation of the current findings concerns the task designs, in which the contingencies in the RDT, intertemporal choice task, and ROVP were set to an ascending order (i.e., increasing delays or probability of punishment). This issue is important as, in some cases, the same manipulation can have opposite effects on choice behavior in block design decision-making tasks such as those used here, depending on whether the contingencies increase or decrease across blocks (Orsini et al, 2018; Orsini et al, 2017b; St. Onge et al, 2010). Such differences in the results of manipulations depending on the order in which choice contingencies are presented have been interpreted as effects on the ability to adapt choices in the context of contingency changes (i.e., behavioral flexibility). As the direction of the effects of reproductive experience was not consistent across tasks (i.e., increases vs. decreases in choice of large, costly rewards), nor were there significant group differences on the probabilistic reversal learning task, it is unlikely that differences in behavioral flexibility fully account for the effects of reproductive experience on decision making. Nevertheless, future studies should address this issue more directly. Another potential limitation concerns possible changes in pain sensitivity in RE females. Although RE and RN females did not differ in shock reactivity, pain is a multi-dimensional construct that is expressed via multiple types of behavior mediated by distinct levels of the neuraxis (Case et al, 2016; Garland, 2012; Melzack & Casey, 1968). Although previous work showed that risk taking in the RDT is unrelated to several measures of pain sensitivity, this work was conducted only in male rats (Simon et al, 2011). It will thus be important in future work to conduct more thorough assessments of pain/shock sensitivity, particularly because parturition is known to cause elevations in pain tolerance that can persist well beyond postpartum (Berlit et al, 2018), and low striatal D2/3 receptor availability predicts high pain threshold in healthy individuals (Martikainen et al, 2018; Pertovaara et al, 2004). Finally, given that performance in both the intertemporal choice task and the RDT is associated with performance in elements of executive function such as working memory and set shifting (Hernandez et al, 2017; Shimp et al, 2015), it would be useful to determine in future work the extent to which effects of reproductive experience on behavior extend to other aspects of PFC-mediated cognition.

### 4.6 Summary and Conclusions

The results of these studies demonstrate a suite of alterations in decision making in RE females (greater risk taking and impulsive choice) that could shed light on post-partum changes in psychiatric disorders, given established links between such psychiatric disorders and both risky and intertemporal choice (Deborah et al, 2020; Johnson et al, 2022; Klein et al, 2008; Pailing & Reniers, 2018; Swann et al, 2008; Takahashi et al, 2008). The results further show that reproductive experience in males (mating) results in an increased preference for delayed over immediate gratification. Additional studies are needed to determine the physiological and neurobiological mechanisms of these effects in both sexes (e.g., via changes in gonadal hormone levels and dopamine signaling), particularly because the vast majority of such work has focused on short postpartum timescales. More importantly, however, given the nearly-ubiquitous prevalence of reproductive experience in humans, these data suggest that reproductive experience should be considered as a variable in preclinical models of human conditions such as aging, in which (at least in women), such experience has been linked to differential risks for brain volumetric changes and even Alzheimer’s disease (Bae et al, 2020; Fox et al, 2018; Jung et al, 2020; Schelbaum et al, 2021).

## Acknowledgements

We thank Vicky Kelley, Brandon Hellbusch, and Bonnie McLaurin for technical assistance. Supported by NIH AG060778 (JLB, BS) and the McKnight Brain Research Foundation (JLB)

## Ethics Statement

The animal study was reviewed and approved by the University of Florida Institutional Animal Care and Use Committee and adhered to the guidelines of the National Institutes of Health.

